# The generation of viable, structurally integrated human-mouse chimaeras through enhanced hPSCs proliferation

**DOI:** 10.1101/2025.06.24.661270

**Authors:** Hideyuki Sato, Ayaka Yanagida, Mariko Kasai, Naoaki Mizuno, Eiji Mizutani, Hiromi Yamamoto, Satoko Ishii, Taro Hihara, Kazuto Yamazaki, Ayuko Uchikura, Kazuaki Nakano, Masashi Ito, Hiroshi Nagashima, Hideki Masaki, Hiromitsu Nakauchi

**Author notes:** Correspondence (H.N.); (A.Y.). These authors contributed equally (Hideyuki Sato, Ayaka Yanagida). Lead contact (A.Y.).

## Abstract

The generation of human organs in animals through blastocyst complementation offers a promising solution to the shortage of transplantable organs. While human pluripotent stem cells (hPSCs) can contribute to interspecies chimaeric embryos when injected into preimplantation embryos of mice, pigs, or monkeys, their integration is often limited due to low chimaerism and segregation from host tissues, significantly impeding progress toward exogenic organ generation. Here, we demonstrate that co-overexpression of the anti-apoptotic gene *BCL2* and the proto-oncogene *MYCL*, along with various cell cycle regulators significantly enhances human cell chimaerism by promoting cell proliferation. This strategy facilitates the generation of viable mouse pups containing hPSC-derived tissues without tumorigenesis. scRNA-seq analysis revealed that hPSCs already exit pluripotent by early post-implantation stage yet hPSCs with enhanced proliferation were able to integrate effectively into the cardiomyocytes and vasculature of both embryonic and extraembryonic tissues with gene expression profiles reflecting their structural integration. These findings highlight the critical role of cell cycle regulation in overcoming xenogeneic barriers. Our findings offer new insights into strategies for enhancing interspecies organogenesis and advancing the field of regenerative medicine.

**Highlight:**

- Enhancing hPSC proliferation increases human cell contribution in interspecies chimaeras
- Achieving the generation of viable human-mouse chimaeras without tumour formation
- hPSCs in post-implantation mouse epiblast exit pluripotency but retain the capacity to integrate into mouse embryogenesis
- hPSCs derivatives integrate into vasculature and cardiac tissues with lineage-matched transcriptional profiles

## INTRODUCTION

Organ transplantation is the only curative treatment for end-stage organ failure. However, it faces significant limitations due to donor organ shortages, immune rejection, and the need for lifelong immunosuppression. Exogenic organ generation through blastocyst complementation, a technique in which pluripotent stem cells (PSCs) are injected into preimplantation embryos genetically modified to lack target organs, offers a promising strategy to generate donor cell-derived organs, even in interspecies animals^1,2^. This approach has been validated in rodent models, including successful organ generation between mice and rats^1,3^. Recently, efforts have been extended to human-mouse interspecies chimaeras^4–7^ and even to human-large animal interspecies chimaeras^7–11^.

Despite these advances, the integration of human cells into host embryos remains inefficient, with human cells often segregating or being eliminated during early embryogenesis^4,12,13^. These challenges highlight the existence of xenogeneic barriers that impede human cell contribution in interspecies chimaeras^14^. Multiple factors have been proposed as potential contributors of xenobarriers, including cell death and insufficient cell-cell adhesion^15^. Gene manipulation techniques, such as the overexpression of anti-apoptotic genes, have been shown to enhance the survival of human PSC (hPSC) derivatives in mouse^5,16–18^ and large-animal embryos^9,10,19^. Additionally, approaches to overcome cell-cell adhesion incompatibility have been tackled to improve human chimaerism^20^. However, the frequency of human cell integration in the host animals remains low, and contributions at later developmental stages have not been thoroughly explored^13^.

To address another potential xenobarrier, we focused on the mismatch in developmental speed between species^21^. Although many cellular and molecular processes are conserved between mice and humans, mouse embryogenesis progresses faster. Several approaches have been explored to mitigate this mismatch, including injecting hPSCs into later-stage host embryos^22^, using lineage-committed donor cells^23^, and introducing growth-promoting oncogenes into hPSCs^16^. However, long-term functional integration of human cells, which is essential for the development of human organs in host animals, has not yet been achieved.

In this study, we investigated the effects of enhanced hPSCs proliferation on chimaerism, differentiation potential and long-term contribution to host embryos. We also examined the capacity of these modified hPSCs to give rise to viable human-mouse chimaeras containing structurally integrated human tissues, without donor-host segregation. Our findings provide new insights into overcoming xenobarriers and advancing interspecies organogenesis.

## RESULTS

### Generation of MYC family gene-overexpressing human-mouse chimaeras

Overexpression of anti-apoptotic genes such as *BCL2* enhanced hPSCs survival in human-mouse chimaera, but the frequency of human cell contribution (chimaerism) remains low^17^. To investigate this limitation, we analysed the gene expression profiles of hPSCs derivatives in embryonic day (E) 6.5 human-mouse chimaeras (see STAR Methods). While the host mouse epiblast remained in a pluripotent state at E6.5, *BCL2*-overexpressing hPSCs (*BCL2*-hPSCs) had exited pluripotency and failed to localise within the epiblast layer (Table S1 and Fig. S1A). To promote retention of pluripotency and support epiblast localisation, we introduced pluripotency factors into *BCL2*-hPSCs. *BCL2*-hPSCs co-overexpressing *OCT4*, *KLF4* and *SOX2* localised to the embryonic region and modestly retained endogenous expression of pluripotency markers at E6.5 (Table S1). However, their derivatives were rarely detected at E9.5. In contrast, *BCL2*-hPSCs co-overexpressing *MYC* (*BCL2*;*MYC*) had also exited the pluripotency by E6.5, but unexpectedly their derivatives persisted in E9.5 embryos (Table S1 and Fig. S1B). This unanticipated contribution led us to expand our investigation to include other members of the MYC family genes: *MYC*, *MYCN* and *MYCL*^24–27^, which are well known to regulate cell proliferation and cell competition across various tissues and species as additional genetic modifications in hPSCs.

We generated hPSC lines constitutively overexpressing *BCL2* and carrying doxycycline (Dox)-inducible *MYC*, *MYCN* or *MYCL* and evaluated their ability to form interspecies chimaeras (Fig. 1A and S1C). As expected, *BCL2*-hPSCs rarely contributed to embryonic tissues or extraembryonic tissues at E13.5 (Fig. 1B), consistent with previous observations at E10.5^17^. In contrast, overexpression of MYC family gene significantly enhanced hPSCs contribution in both embryonic and extraembryonic regions. A previous study showed that *BCL2* and *MYCN*-overexpressing hPSCs were present in E10.5 mouse embryos^16^. However, our data revealed that hPSCs overexpressing *BCL2* and either *MYC* or *MYCN* (*BCL2*;*MYC* or *BCL2*;*MYCN*) formed disorganised cell masses and failed to structurally integrate into host mouse tissues at E13.5. Bulk RNA-sequencing (RNA-seq) of cell masses isolated from the mediastinum and intestines of E13.5 *BCL2*;*MYC*-hPSC human-mouse chimaeras showed differentiation into multiple lineages (Fig. S1E). In contrast, hPSCs overexpressing *BCL2* and *MYCL* (*BCL2*;*MYCL*) did not form such disorganised masses and instead contributed to both embryonic and extraembryonic tissues, forming organised tissue structures.

**Figure 1.**
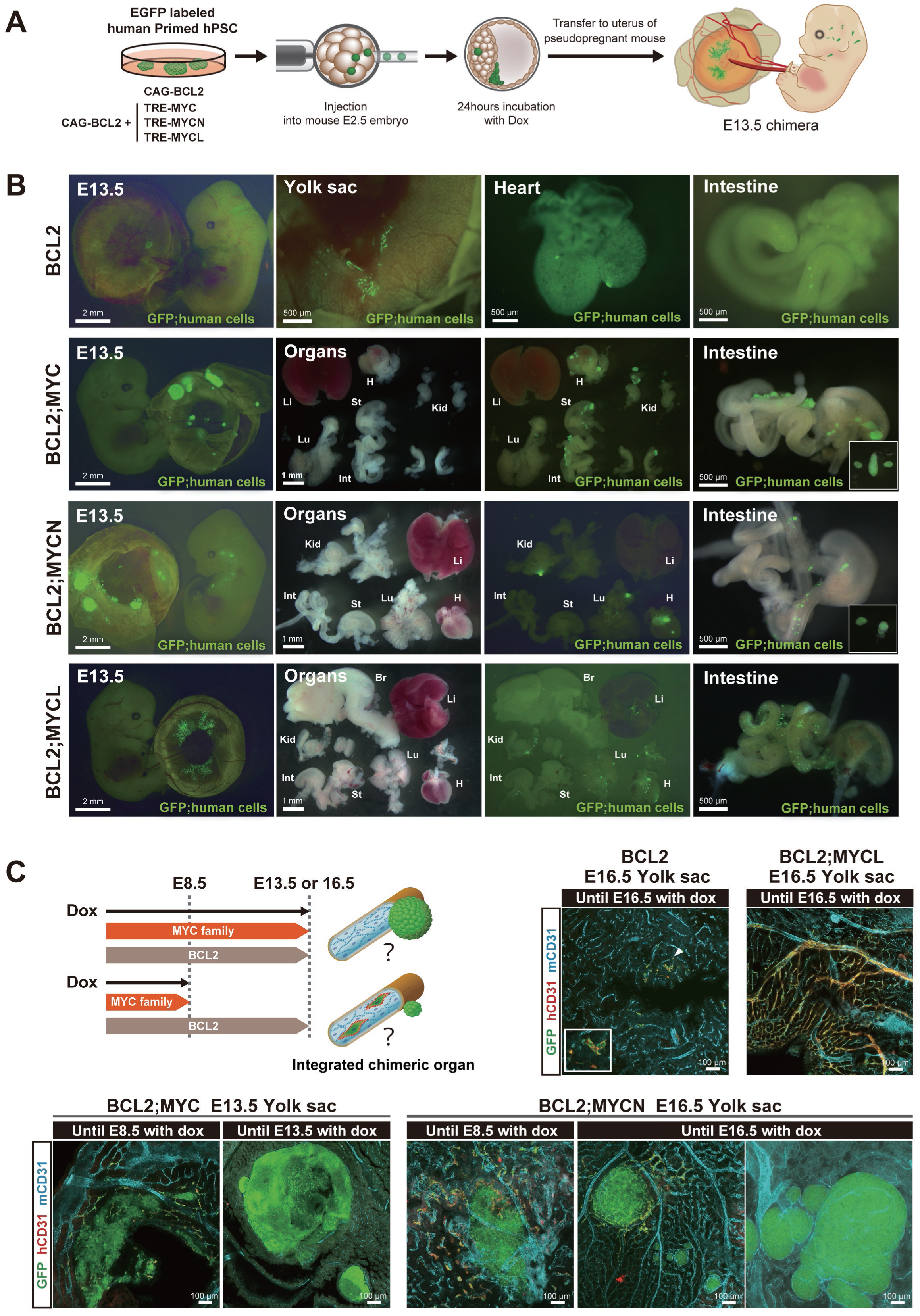
Analysis of human cell contribution in human-mouse chimaeras using hPSCs overexpressing *BCL2* and *MYC* family genes. (A) Schematic overview of the generation of human-mouse chimaeras. (B) Representative images of E13.5 human-mouse chimaeras generated derived from GFP-labelled hPSCs overexpressing *BCL2* alone (BCL2) or in combination with *MYC*, *MYCN* or *MYCL* (BCL2;MYC, BCL2;MYCN, BCL2;MYCL). Insets show isolated GFP-positive human cell clusters within the intestines. Li, liver; Lu, lung; St, stomach; H, heart; Int, intestine; Kid, kidney. (C) Schematic of Dox administration timelines and representative images of yolk sac from E13.5 or E16.5 chimaeras derived from GFP-labelled *BCL2*-, *BCL2*;*MYC*-, *BCL2*;*MYCN*- or *BCL2*;*MYCL*-hPSCs under varying duration of Dox administration. Insets show high-magnification images of the region indicated by an arrowhead. mCD31, mouse CD31; hCD31, human CD31. Single-channel images are shown in Fig. S1D. See related Figure S1.

### Duration of Myc family gene induction regulates the structural contribution of hPSCs to human-mouse chimaeras

To investigate whether the duration of MYC family gene induction affects tumorigenesis and the differentiation potential of hPSCs in human-mouse chimaeras, we examined the effects of Dox administration duration (Fig.1C and S1C). In *BCL2*;*MYC*- or *BCL2*;*MYCN*-hPSC chimaeras, limiting Dox administration to E8.5 resulted in hPSC-derived cell masses surrounded by endothelial cell marker^28^ CD31^+^ hPSC-derived vascular structures in the yolk sac at E13.5 or E16.5. However, extending Dox administration to E13.5 or E16.5 led to increased hPSC-derived cell mass size, while the proportion of hPSC-derived vascular structures significantly decreased (Fig. S1D). *BCL2*;*MYCN*-hPSCs also formed cardiomyocyte marker ACTC1^29^ negative cell masses in the heart (Fig. S1F). In contrast, prolonged Dox administration in *BCL2*-alone- or *BCL2;MYCL*-hPSC chimaeras up to E16.5 did not lead to the formation of hPSC-derived cell mass. Instead, *BCL2;MYCL*-hPSCs contributed to well-organised CD31⁺ vascular networks in the yolk sac. These findings suggest that the duration of MYC family gene induction critically influences the balance between structured tissue integration and disorganised cell proliferation, with *MYCL* supporting sustained, tumour-free contribution even under prolonged induction conditions.

### Accelerating cell proliferation enhances hPSC chimaerism in human-mouse chimaeras

MYC family genes are well known to promote cell proliferation^30,31^. We, therefore, investigated whether enhancing cell proliferation could improve the chimaeric potential of hPSCs in human-mouse chimaeras (Fig. 2A). To this end, we generated hPSC lines constitutively expressing *BCL2* and carrying Dox-inducible transgenes encoding cell-cycle accelerators–*CDK9*, *E2F1*, *E2F2*, *E2F3a* or *E2F4*^32,33^. Upon Dox induction, all lines overexpressing MYC family genes or cell-cycle accelerators except E2F1 showed enhanced proliferation in vitro (Fig. 2B). Consistent with previous reports describing *E2F1*’s unique pro-apoptotic activity among E2F family members^34–37^, E2F1 overexpression instead induced cell death. We next evaluated these modified lines’ in vivo chimaera-formation capacity. The rates of total live foetuses and chimaeric foetuses were not significantly different between highly proliferative hPSC lines (Fig. 2C and S2A).

**Figure 2.**
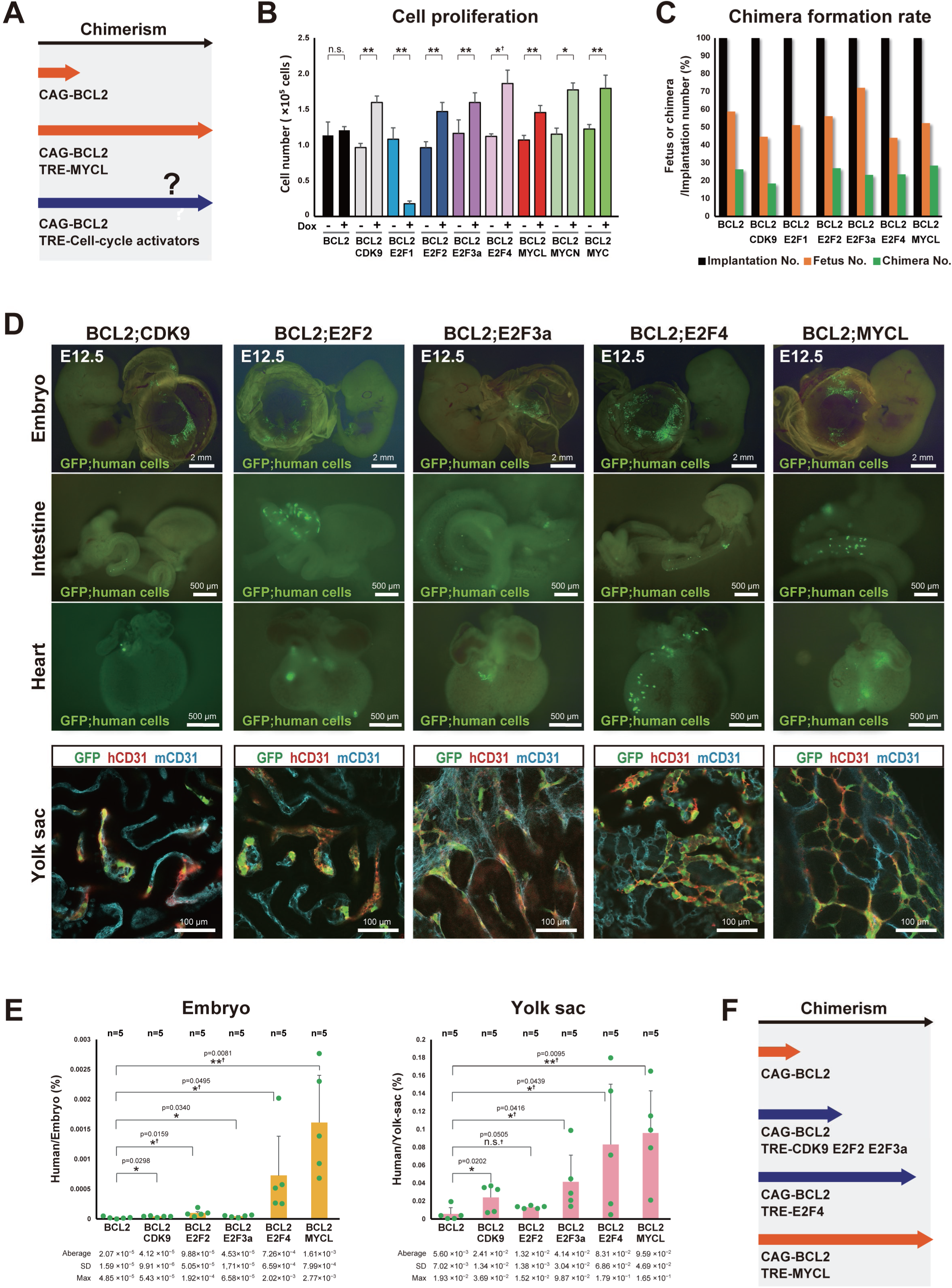
Accelerating cell proliferation enhances hPSC chimaerism in interspecies chimaeras. (A) Schematic model illustrating the hypothesis of relationship between enhanced cell proliferation and increased human cell chimaerism. (B) Cell counts of hPSCs overexpressing *BCL2* alone or in combination with a cell cycle-related gene (*CDK9*, *E2F1*, *E2F2*, *E2F3a*, or *E2F4*), *MYC*, *MYCN* or *MYCL*, with or without Dox treatment over a single passage (n=3 each). (C) Chimaera formation rates at E12.5 human-mouse chimaeras. Dox was administered until the sampling time point. (See also Fig. S2A) (D) Representative images of whole embryos, intestines, hearts and yolk sacs from E12.5 human-mouse chimaeras. mCD31, mouse CD31; hCD31, human CD31. (E) ddPCR analysis of human cell chimaerism in whole embryos and yolk sac from E12.5 chimaeras. Each dot represents one chimaeric embryos. (F) Schematic model illustrating the relationship between enhanced cell proliferation and increased human cell chimaerism. Graphs show mean ± standard deviation (SD). Statistical significance was assessed using unpaired Student’s t-test, except for comparisons marked with †, which were analyzsed using Welch’s t-test. n.s., not statistically significant; *p < 0.05, **p < 0.01. See related Figure S2.

We further analysed the spatial contribution of hPSC derivatives in E12.5 chimaeric embryos (Fig. 2D). hPSCs overexpressing cell-cycle accelerators contributed to both embryonic tissues (e.g., intestine and heart) and extraembryonic tissues (yolk sac), similar to the contribution pattern of *BCL2*;*MYCL*-hPSCs. In the yolk sac, hPSC-derived cells were structurally integrated into the vasculature, expressed CD31, and connected with host-derived blood vessels. To quantify human cell chimaerism, we performed droplet digital PCR (ddPCR) targeting a human mitochondrial DNA sequence in E12.5 embryonic and yolk sac tissues. *BCL2*-hPSCs co-overexpressing *CDK9*, *E2F2*, *E2F3a*, *E2F4,* or *MYCL* exhibited higher levels of chimaerism (0. 04–1.6 × 10^−3^% in embryos and 1.3–9.59 × 10^−2^% in yolk sacs) compared with *BCL2*-alone hPSCs (0.02 × 10^−3^% in embryos; 0.56 × 10^−2^% in yolk sacs) (Fig. 2E). However, the degree of chimaerism enhancement varied across the cell-cycle accelerators, with *E2F4* and particularly *MYCL* achieving the significantly highest levels (Fig. 2F). We also assessed chimaera formation efficiency and chimaerism using another *BCL2*;*MYCL*-hPSC line (7F3669#1), which exhibited similar chimaeric embryo rates (26–29%) (Fig. S2B), spatial contribution patterns (Fig. S2C) and chimaerism levels (Fig. S2D). Based on these results, we conducted further analysis of human-mouse chimaeras derived from *BCL2*;*MYCL*-hPSCs.

### Cell lineages and function of *BCL2*;*MYCL*-hPSC derivatives in human-mouse chimaeras

We next examined whether *BCL2*;*MYCL*-hPSCs differentiated into cell lineages that contribute to host tissues in human-mouse chimaeras. Detailed dissection confirmed that *BCL2*;*MYCL*-hPSC derivatives contributed to E12.5 yolk sac and various embryonic tissues, including internal organs such as the liver, diaphragm, heart and intestine, as well as skin (Fig. 3A and S3A). Among these, the heart exhibited the highest level of contribution, with 3.30 × 10^−2^% chimaerism (Fig. S3B). To characterise the cell lineage of *BCL2*;*MYCL*-hPSC derivatives, we performed single-cell RNA sequencing (scRNA-seq) on *BCL2*;*MYCL*-hPSCs and their derivatives isolated from E12.5 chimaeric hearts and intestines, which consistently exhibited the highest levels of contribution across the chimaeric embryos. These derivatives did not express significant levels of pluripotent stem cell markers such as *POU5F1* (*OCT4*)^38,39^, *NANOG* ^40,41^, SOX2^42,43^ and *CDH1*^44^(Fig. 3B). Instead, *BCL2*;*MYCL*-hPSC derivatives isolated from the heart expressed cardiomyocyte markers such as α*-cardiac actin* (*ACTC1*)^29^*, tropomyosin* α*-1* (*TPM1*)^45^, and troponin T (*TNNT2*)^46^, whereas derivatives from the intestine upregulated vascular endothelial markers including *CD31* (*PECAM1*)^28^*, KDR* (*VEGFR2*)^47^ and *CDH5* (*VE-cadherin*)^48^. To exclude the possibility that these cells arose from cell fusion of human and mouse cell^49,50^, we performed genomic PCR targeting human and mouse β-actin genes on GFP-positive and GFP-negative fractions sorted by flow cytometry from E12.5 chimaeric hearts. GFP-positive cells contained only human genomic DNA, confirming that they were not the result of cell fusion (Fig. S3C and S3D).

**Figure 3.**
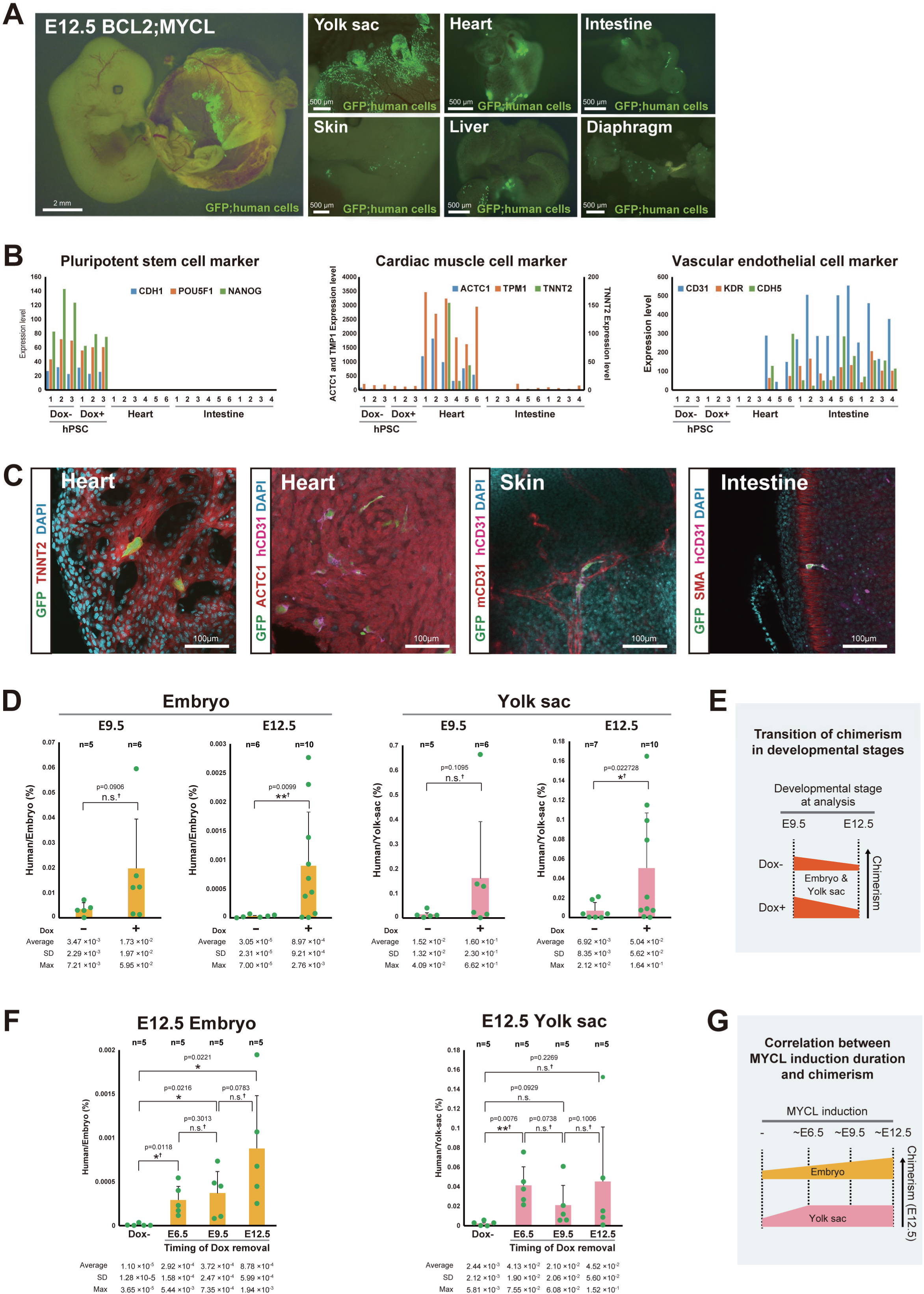
Localisation, identity and chimaerism of hPSC-derived human cell in human-mouse chimaeras. (A) Representative images of E12.5 human-mouse chimaeras derived from GFP-labelled *BCL2*;*MYCL*-hPSCs. Dox was administered until the time of dissection. (B) scRNA-seq analysis of gene expression profiles in hPSCs and human cells isolated from hearts and intestines of E12.5 human-mouse chimaeras generated from *BCL2*;*MYCL*-hPSCs. (C) Representative images of hearts, skin and intestines from E12.5 human-mouse chimaeras derived from GFP-labelled *BCL2*;*MYCL*-hPSCs. (D) ddPCR analysis of human cell chimaerism in whole embryos, isolated yolk sacs from E9.5 or E12.5 human-mouse chimaeras derived from *BCL2*;*MYCL*-hPSCs. Each dot represents one embryo or tissue. (E) Schematic summary of human cell chimaerism dynamics across developmental stages. (F) ddPCR analysis of human cell chimaerism in whole embryos and isolated yolk sacs from a single E12.5 human-mouse chimaeras derived from *BCL2*;*MYCL*-hPSCs. Dox was administered until the developmental stage indicated. Each dot represents one embryo or tissue. (G) Schematic model illustrating the relationship between the duration of cell cycle accelerator induction and the extent of human cell chimaerism in human-mouse chimaeras. Graphs show mean ± standard deviation (SD). Statistical significance was assessed using unpaired Student’s t-test, except for comparisons marked with †, which were analyzed using Welch’s t-test. n.s., not statistically significant; *p < 0.05, **p < 0.01. See related Figure S3.

We also investigated the localisation and lineage identity of *BCL2*;*MYCL*-hPSCs derivatives by immunofluorescent staining. In E12.5 human-mouse chimaera, *BCL2*;*MYCL*-hPSC derivatives within the heart expressed TNNT2 or CD31 (Fig. 3C). In the skin, *BCL2*;*MYCL*-hPSC derivatives also expressed CD31 and formed vascular networks integrated with the host mouse vasculatures. In the intestines, *BCL2*;*MYCL*-hPSC derivatives were detected in non-endodermal layers, particularly in the α-smooth muscle actin (SMA)-positive mesodermal wall as CD31^+^SMA^-^ cells. To assess functionality, we examined *BCL2*;*MYCL*-hPSCs derivatives isolated from E13.5 human-mouse chimaeric hearts. These cells exhibited spontaneous beating in vitro (Video S1). These results demonstrate that hPSC derivatives localise appropriately within host mouse tissues and differentiate into lineages-appropriate and functional cell types.

### Prolonged induction of cell-cycle accelerator attenuates the decline in hPSC chimaerism during embryogenesis

To determine whether the timing and duration of proliferation enhancement influence the extent of human cell contribution, we examined hPSC chimaerism across multiple developmental stages (Fig S3A). ddPCR analysis revealed that in chimaeras without Dox treatment, hPSC chimaeras at E12.5 declined to approximately 0.9% of the levels observed at E9.5. In contrast, prolonged Dox administration to induce a cell cycle accelerator attenuated this decline (Fig. 3D-E and S3E). A similar trend was observed in the yolk sac.

To further define the temporal window during which enhanced proliferation supports robust chimaerism, we varied the duration of Dox administration and analysed chimaerism at E12.5. In embryonic tissues, the lowest levels of chimaerism were observed in chimaeras with Dox withdrawal at E6.5, whereas continued induction until E12.5 resulted in significantly higher chimaerism (Fig. 3F). In contrast, chimaerism in the yolk sac was largely unaffected by the duration of Dox treatment, suggesting that the proliferative effect of cell-cycle accelerators and the requirement for such stimulation may vary depending on the target tissue (Fig. 3G).

### Birth of structurally integrated human-mouse chimaeras

To determine whether enhancement of hPSC proliferation enables persistent integration into functional tissues, we evaluated human–mouse chimaeras at E19.5. Specifically, we examined chimaeras derived from *BCL2*;*MYCL*-hPSCs and hPSCs co-expressing Dox inducible *MYCL* and an alternative anti-apoptotic gene, *BCL2L1* (*BCL2L1*;*MYCL*-hPSCs) (Fig. S4A). Despite Dox withdrawal at E9.5, *BCL2L1*;*MYCL*-hPSC derivatives contributed robustly to both embryonic tissues (heart, intestine, and portal vein) and extraembryonic tissues (yolk sac) (Fig. S4B). Similarly, continuous Dox administration up to E19.5 allowed *BCL2*;*MYCL*-hPSCs to persist in both embryonic and extraembryonic tissues (Fig. 4A), consistent with their contribution at earlier developmental stages (Fig. 1B and 3A).

**Figure 4.**
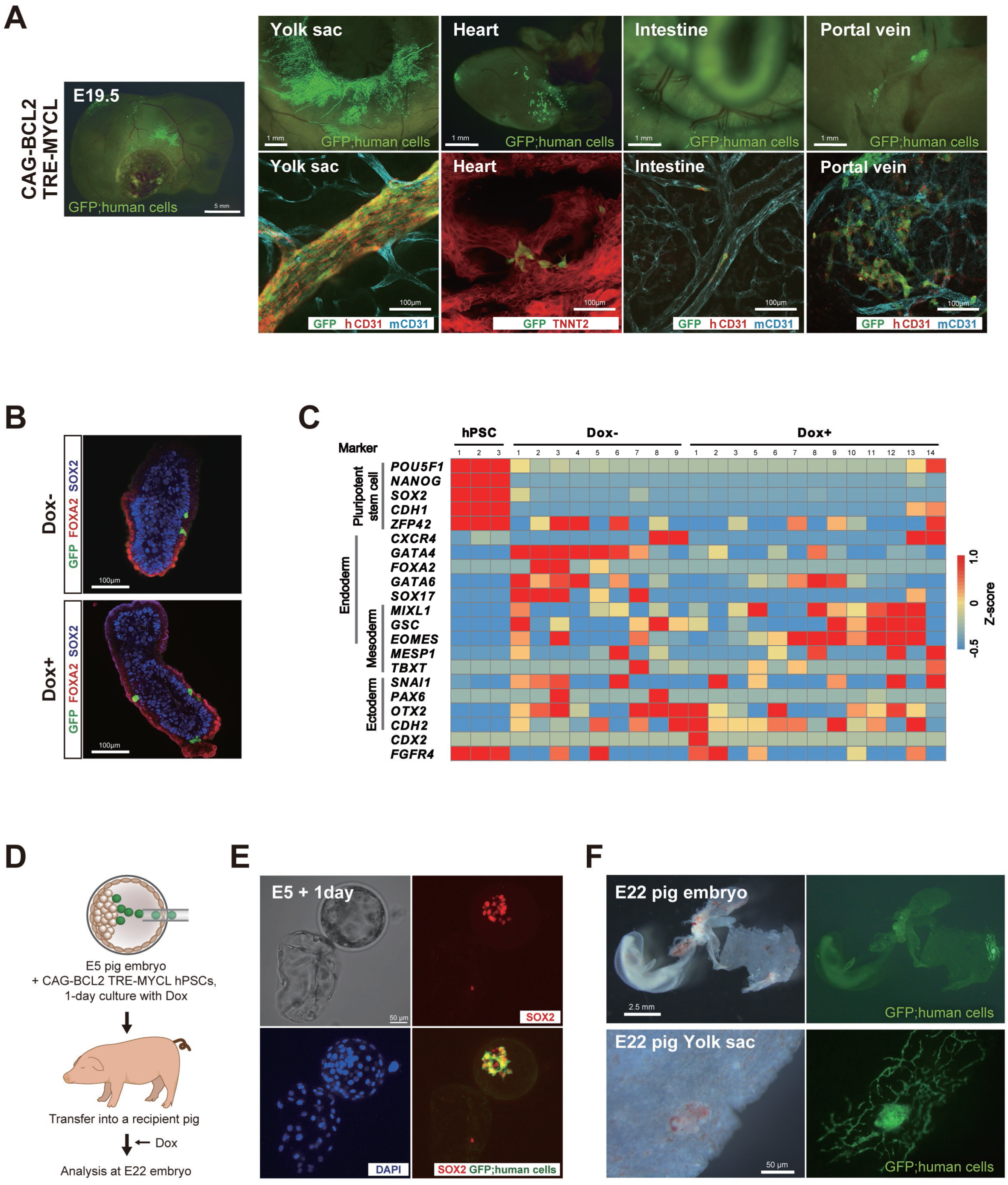
Developmental progression and lineage integration of hPSC-derivatives in human-mouse and human-pig chimaeras. (A) Representative images of E19.5 human-mouse chimaeras derived from GFP-labelled *BCL2*;*MYC*-hPSCs. Dox was administered until the time of dissection. (B) Representative images of E6.5 human-mouse chimaeras derived from GFP-labelled *BCL2*;*MYCL*-hPSCs. (C) Heatmap of gene expression profiles from scRNA-seq of undifferentiated hPSCs and *BCL2*;*MYCL*-hPSC derivatives isolated from E6.5 human-mouse chimaeras. (D) Schematic overview of the generation protocol for human–pig chimaeras. (E) Representative images of E6 pig embryos injected with GFP-labelled *BCL2*;*MYCL*-hPSCs, followed by one day of in vitro culture with Dox. SOX2, DAPI and merge images are shown as maximum intensity projection. (F) Representative images of E22 pig embryos injected with GFP-labelled *BCL2*;*MYCL*-hPSCs. Bottom panels show high-magnification images of the yolk sac. See related Figure S4.

Notably, hPSC derivatives formed structured CD31^+^ vascular networks in the yolk sac, TNNT2^+^ cardiomyocytes in the heart, and CD31^+^ endothelial cells within the intestine and portal vein without forming tumour-like cell masses (Fig. 4A). In contrast, their contribution to the brain was limited, with an average chimaerism of only 3.42 × 10^−5^% (n=4) (Fig. S4C). These results demonstrate that promoting hPSCs proliferation enables not only prolonged survival and contribution but also the generation of viable live human-mouse chimaeras with structurally integrated human cells within cardio-vasculature tissues.

### Developmental trajectory of hPSCs in early human-mouse chimaeras

To determine how and when lineage-biased contributions of hPSCs emerge during development, we analysed the differentiation dynamics of *BCL2*;*MYCL*-hPSC derivatives in early-stage human-mouse chimaeras. At E3.5, derivatives of both *BCL2*-hPSCs with and without *MCYL* induction were partially localised to the inner cell mass (ICM) region of host blastocysts (Fig. S4E). By E6.5, *MYCL* induction increased contribution of human cells compared to embryos without induction (Fig. 4B and S4F). However, even with *MYCL* induction, the hPSC derivatives were dispersed throughout the embryo and were primarily localised at the interfaces between the epiblast and extra-embryonic ectoderm (ExE), between the visceral endoderm and epiblast, and within both EXE and epiblast.

To characterise the lineages identities of these cells, we performed scRNA-seq on *BCL2*;*MYCL*-hPSC derivatives isolated from the E6.5 embryonic region. These cells showed downregulation of pluripotent stem cell markers (*NANOG*, *OCT4*, *SOX2,* and *CDH1*) and upregulation of lineages-specific markers representing all three germ layers (Fig. 4C). In embryos without *MYCL* induction, hPSC derivatives predominantly expressed endodermal markers such as *GATA4*, *GATA6* and *SOX17* although subsets expressed mesodermal or ectodermal markers. In contrast, in embryos with *BCL2* and *MYCL* induction, most hPSC derivatives expressed markers characteristic of mes-endodermal precursors and mesodermal lineages, including *MIXL1*, *GSC*, and *EOMES*, while expression of the mesodermal marker *TBXT* (Brachyury) remained low. Although further studies are required to precisely define the lineage identities of these cells and map them to corresponding stages of human embryogenesis, our findings suggest that hPSCs derivatives undergo early multi-lineage differentiation by E6.5. Moreover, enhanced cell cycle acceleration appears to influence lineage specification, potentially contribution to increased integration into cardiovascular tissues at later developmental stages.

### hPSCs contribute to vascular-like structures of pig embryos

To assess whether the integrative potential of proliferation-enhanced hPSCs extends beyond mice hosts, we examined their contribution in pig embryos. We injected 10–20 *BCL2*;*MYCL*-hPSCs into parthenogenetic pig blastocysts at E5 (Fig. 4D). Within one day, the majority of injected cells localised to the ICM region, regardless of *MYCL* induction via Dox treatment, and retained high expression of the core pluripotency marker SOX2 (Fig. 4E). To evaluate hPSCs’ contribution in post-implantation embryos, the injected blastocysts were transferred into the uteri of oestrus-synchronized recipient gilts, and embryos were collected at E22. *BCL2*;*MYCL*-hPSCs derivatives were detected in the yolk sac, where they contributed to vasculature-like branched structures (Fig. 4F and S4G). These results demonstrate that co-overexpression of *BCL2* and *MYCL* enhances the ability of hPSCs to survive and structurally integrate into the tissues of large-animal host. This cross-species contribution highlights the broader applicability of this stagey for advancing interspecies chimaerism and supports its potential in future efforts toward exogenic organ generation.

## DISCUSSION

In this study, we demonstrate that enhancing donor hPSC proliferation significantly improves human cell chimaerism, enables the formation of structurally integrated and functional tissues without tumorigenesis, and leads to the birth of viable human-mouse chimaeras. These findings represent an important step toward the advancement of exogenic organ generation through blastocyst complementation.

Previous studies using *BCL2*-hPSCs reported limited chimaerism and lacked vascular contribution^16,17^, unlike the vascular integration observed in our study (Fig.1B and S1D). Our use of multiple hPSC lines (Fig. S2B-D) suggests that these differences are unlikely to clone-specific variability. Instead, culture conditions such as the use of the Tankyrase inhibitor XAV939 or different basal media may impact hPSCs heterogeneity and differentiation potential^52,53^.

To identify a gene that supports robust and sustained hPSC integration, we compared MYC family members and found that *MYCL* uniquely enabled long-term human cell contribution without tumorigenesis. While *MYCN* has previously been shown to enhance human cell chimaerism at E10.5, this effect required Flk1-haploinsufficient mouse hosts and did not assess longer-term integration outsomes^16^. In our study, prolonged *MYCN* induction resulted in disorganised cell masses and segregation from host tissues (Fig.1B-C and S1D). In contrast, *MYCL*, which has comparatively lower tumorigenic activity^54–56^, makes it a safer and more effective candidate for exogenic organ generation.

We show that accelerating the cell cycle in hPSCs enhances human cell chimaerism without altering the preferential localisation within host tissues. Given the more rapid pace of mouse embryogenesis compared to human development, this temporal mismatch likely constitutes a key xenobarrier. Our data suggest that promoting hPSC proliferation may partially mitigate this developmental pace mismatch. Notably, *BCL2*;*E2F4*- and *BCL2*;*MYCL*-hPSCs achieved the highest chimaerism levels among all tested lines. These differences may reflect gene-specific mechanisms of action, with cell cycle regulators influencing proliferation in stage- and tissue-specific ways in vivo. Despite these improvements, mouse host cells may still differentiate more rapidly than donor human cells, potentially contributing to the observed decline in chimaerism at later developmental stages (Fig. 3D-E). Our previously reported “cell-competitive niche” strategy^57^ may further mitigate this mismatch and promote sustained human cell integration.

While enhanced proliferation improved overall chimaerism, the majority of hPSCs were still segregated from the post-implantation epiblast layer by E6.5 (Fig. 4B and S4F), consistent with earlier studies^4,58^. However, our results show that hPSC derivatives can contribute to spatially appropriate, structurally organised tissues (Fig. 4A and S4B). This suggests that enhanced proliferation alone is insufficient to overcome the peri-implantation xenobarriers, particularly segregation from the host epiblast, However, a subset of hPSC derivatives can reintegrate into host embryogenesis during gastrulation.

We observed that hPSCs preferentially contributed to cardio-vascular tissues in interspecies chimaeras (Fig. 1B and 2D). Vasculature integration of hPSCs has been reported in human-mouse^16^ and human-pig chimaeras^9^ using *Flk*- or *ETV2*-deficient hosts, whereas robust contribution to solid organs remains a major challenge. This biased contribution may be attributable to early lineage specification bias observed in E6.5 *BCL2*;*MYCL*-hPSC derivatives (Fig. 4C), as well as to a degree of lineage-specific permissiveness in the interspecies context, potentially due to conserved developmental cues. In contrast, endodermal contribution was limited, suggesting selective elimination by additional xenobarriers. In mouse-mouse chimaeras, forced expression of *MIXL1* has been shown to enhance endodermal contribution^60^, raising the possibility that targeted lineage priming may improve the generation of endoderm-derived organs in interspecies chimaeras.

Given that hPSC derivatives contributed to both yolk sac and embryonic vasculature, as well as to cardiomyocytes, these cells may contribute to embryonic and extraembryonic-mesodermal lineages during gastrulation. Although the origin of human extraembryonic mesoderm remains debated^59^, if hPSCs indeed differentiated into these lineages, interspecies chimaera may provide a valuable model for studying hPSCs differentiation potential and early human embryogenesis in vivo.

### Limitations of this study

The precise dynamics and routes of human cell contribution during early post-implantation development remain incompletely understood. Future studies employing high-resolution time-lapse imaging and improved ex vivo post-implantation embryo culture systems with high efficiency will be required to track human cell behaviour in interspecies chimaeras and to define the developmental windows during which mismatches in lineage identity and spatial localisation can be buffered in human-mouse chimaeras. Moreover, we did not detect appreciable human cell contribution to the embryonic region in pig chimaeras. This may reflect species-specific developmental incompatibilities or currently unidentified other xenobarriers. Investigations using additional large-animal models, including non-human primates, will be necessary to assess the broader applicability of this approach to human organ generation. Finally, although human cells contributed to cardiomyocytes and yolk sac vasculature in human-mouse chimaeras, the corresponding stages of human foetal development remain undefined. A recent study in rat-mouse chimaeras demonstrated that rat-derived organs did not fully mature within the shorter developmental timeline of mouse hosts^61^. Similarly, in human-mouse chimaeras, differences in developmental pace between species and potential incompatibilities in ligand-receptor interactions may influence the maturation of contributed human tissues, while the kinetics of hPSCs differentiation in xenogeneic environments remains debated^62,63^. Further development of high-resolution, stage-specific reference atlases of human embryogenesis will be required to accurately map the developmental status of human cells in interspecies chimaeras.

## Acknowledgements

We are grateful to Toshihiro Kobayashi for providing critical feedback, Motoo Watanabe, Kyoko Okada and Hiroko Tsukui for laboratory assistance. We would like to acknowledge support staff in Stem Cell Therapy Laboratory and animal facility of Institute of Science Tokyo, and members of the Nakauchi Lab at Stanford University for discussion and comments. This research was supported by AMED (JP22bm1004002 and JP23bm1123041 to H.N.), the Leducq Foundation and Ludwig Foundation (H.N.) JST FOREST Program (JPMJFR214Y to A.Y., JPMJFR235J to N.M.) and JSPS KAKENHI (22K07886 to A.Y., 20H03170, 23H02401 to H.M., 21K07837 to N.M.).

## Author contribution

Conceptualisation, HS, HN; Methodology, HS, AY; Investigation; HS, AY, MK, NM, EM, SI, TH, KY; Formal analysis, HS, NM, SI, TH, KY; Resources, AU, KN, HN; Funding acquisition, AY, NM, HM, HN; Writing–original draft, AY, HS; Writing–revision and editing, AY, HS, NH, NM, SI, TH, KY, HM; Supervision, AY, MI, HN, HM, HN

## Declaration of generative AI and AI-assisted technologies in the writing process

During the preparation of this work, the authors used ChatGPT (OpenAI) to improve grammar of the text. After using this tool, the authors carefully reviewed and edited the content to ensure accuracy and appropriateness, and take full responsibility for the content of the publication.

## Materials and Methods

### Experimental model, subject details with ethics information

#### Human samples

The human primed iPSC lines 7F3669#1, PB001 and PB004 (Masaki et al., 2015) were derived from neonatal human dermal fibroblast (Lonza) and peripheral blood samples obtained from volunteers after written informed consent, respectively, by reprogramming using a Sendai virus vector (Nishimura et al. 2011) expressing human *OCT4*, *SOX2*, *KLF4* and *MYC*. The collection of peripheral blood samples and the generation of iPSC were approved by the ethical committee of the Institute of Tokyo Science and the University of Tokyo. Research involving human-mouse interspecies chimaera was approved by Research Ethics Committee of the University of Tokyo and the Ministry of Education, Culture, Sports, Science and Technology Japan following confirmation of compliance by the Specified Embryo Expert Committee. Research involving human-pig interspecies chimaera was approved by the Research Ethics Committees of the University of Tokyo and Meiji University, and by MEXT Japan following confirmation of compliance by the Specified Embryo Expert Committee.

#### Mouse

ICR, BDF1 and C57BL/6N mice were obtained from CLEA Japan and maintained in a biofacility with daily health checks carried out by dedicated trained staff. The mice were maintained on a lighting regime of 12:12 hours light:dark with food and water supplied ad libitum. Use of animals in this project was approved by the animal committee for Institute of Science Tokyo.

#### Mouse embryos

BDF1 or ICR female mice were superovulated by intraperitoneal injection with 5 IU of PMSG, followed by an injection of 5 IU of hCG 48 hour (h) later, and subsequently intercrossed with C57BL/6N or ICR males. 8-cell/morulae stage embryos were collected with M2 medium from the oviduct of the female mice at 2.5 days post-coitum (dpc). The embryos were transferred into KSOM medium covered with mineral oil and cultured at 37°C in 5% CO_2_. Embryos at the 8-cell/morulae stages were cryopreserved in cryovials containing DAP213 medium and stored in liquid nitrogen until use. Cryopreserved 8-cell/morulae-stages embryos were thawed 2-3 h before injection using 0.25 M sucrose solution and cultured in KSOM until injection.

#### Pig and pig embryos

Pig ovaries were collected at a local abattoir and transported to the laboratory in DPBS containing 75 µg/mL potassium penicillin G, 50 µg/mL streptomycin sulfate, 2.5 µg/mL amphotericin B and 0.1% (w/v) polyvinyl alcohol. Cumulus-oocyte complexes (COCs) were collected by aspiration from ovarian antral follicles with diameters of 3.0-6.0 mm. COCs with at least three layers of compacted cumulus cells were selected and cultured in POE-CM (Research Institute for the Functional Peptides). The COCs were cultured for 22 hours with eCG and hCG in a humidified atmosphere of 5% CO_2_ and 95% air at 38.5°C. HP-POM (Research Institute for the Functional Peptides) supplemented with 50 ng/mL rhFSH (RayBiotech) and 1 mM dibutyryl cAMP (dbcAMP, Research Institute for the Functional Peptides) was used for the next 20 hour in a humidified atmosphere of 5% CO_2_ and 5% O_2_ at 38.5°C. The COCs were then cultured for 22–24 hours without rhFSH and dbcAMP under the same conditions. In vitro matured oocytes with expanded cumulus cells were treated with 1 mg/mL hyaluronidase dissolved in Tyrode’s lactose medium containing 10 mM Hepes and 0.3% (w/v) polyvinylpyrrolidone (Hepes-TL-PVP) and separated from the cumulus cells by gentle pipetting. Oocytes with an evenly granulated ooplasm and an extruded first polar body were selected for subsequent experiments. Parthenogenetic zygotes were cultured in PZM-5 medium (Research Institute for the Functional Peptides) under paraffin oil in a humidified incubator at 38.5°C in 5% CO_2_ and 5% O_2_. Embryos beyond the morulae stages were cultured in PZM-5 medium supplemented with 10% FBS. Crossbred (Large White/Landrace x Duroc) prepubertal gilts, weighing from 105 kg, were used as recipient of human-pig interspecies embryos. All procedures involving pigs were approved by the Institutional Animal Care and Use Committee and Research Ethic Committee of Meiji University.

#### Cell Culture

Cell lines were cultured in humidified incubators at 37°C in 5% CO_2_ and 5% O_2_ unless specified. They are confirmed negative for mycoplasma by periodic screening.

### Method details Cell culture

hPSCs were propagated in a bFGF-based medium (StemFlex) supplemented with 10 µM of XAV939, a WNT inhibitor, on Matrigel-coated dishes. 2 µM Rock inhibitor (Y-27632 [Y]) was added to the media during replating. Cells were passaged by dissociation with 0.25% Trypsin-EDTA every 3-4 days.

Inactivated Mouse embryonic fibroblasts (MEFs) were cultured in DMEM-high glucose medium supplemented with 5% and 1% penicillin-streptomycin-glutamine (PSG) at 37°C in 5% CO_2_. 293T cell lines were cultured in DMEM-high glucose medium supplemented with 10% FBS and 1% PSG at 37°C in 10% CO_2_.

### Lentiviral vector production

A human *BCL2* and *BCL-XL* expressing lentiviral vector plasmids were constructed from CS-CAG-GFP. Dox-inducible human MYC family gene (*MYC*, *MYCN*, *MYCL*), cell cycle regulatory gene (*E2F1*, *E2F2*, *E2F3a*, *E2F4*, *CDK9*), and *BCL-XL* lentiviral vectors plasmid were derived from CS-TRE-PRE-Ubc-rtTA-IRES2-EGFP or CS-TRE3G-PRE-Ubc-rtTA3G-IRES-EGFP (Yamaguchi et al., 2012). The open reading frames (ORFs) of the genes above were amplified from cDNA synthesized from total RNA extracted from human primed PSCs (PB001). Each of these ORFs was inserted into the lentiviral vector immediately downstream of the TRE, and a puromycin, neomycin, hygromycin, blasticidin, or Zeocin resistance gene was inserted immediately downstream of IRES, using In-Fusion Snap Assembly Master Mix. Lentiviruses were produced by co-transfection of the lentiviral vector plasmid with packaging plasmids (pMDLg/pPRE and pCMV-VSV-G-RSV-Rev). 293T cells were seeded at 3.5×10^6^ cells per 10 cm dish one day before transfection. Calcium phosphate transfection was used to transfect the 293T cells with 15 µg lentiviral vector and 10 µg of each packaging plasmid. The culture medium was exchanged 16 h after transfection, and the cells were incubated for 48 h. Lentiviruses were concentrated by ultracentrifugation at 4000 x *g* for 2 h, and the pellet was resuspended in 1/250th volume of PBS. Concentrated viruses were stored at –80°C.

#### Generation of transgenic hPSCs

Transgenes were introduced into hPSCs by lentiviral transduction or safe-harbor gene knock-in. Lentiviral transduction were performed by co-culture of feeder-free hPSCs and the viruses for 1-2 days. Puromycin (500 ng/mL), G418 (800 µg/mL), hygromycin (200 µg/mL), blasticidin (5 µg/mL) or Zeocin (5 µg/mL)-resistant clones were selected after 3 days of lentivirus transduction, and GFP-expressing hPSCs were purified by FACS sorting and propagated. In the case of safe-harbor knock-in, 3^rd^ generation TetOn-expression cassette for 2A-linked codon-optimized human OKS (*POU5F1*, *KLF4*, *SOX2*) and codon-optimized human *BCL2* followed by CAG-tdTomato-2A-TetOn3G was inserted into *AAVS1* locus by CRISPR-Cas9 genome editing. The targeting vector also contains puromycin resistant cassette. Knock-in clones were enriched following FACS-sorting of tdTomato+ cells following 250 ng/mL puromycin selection and further confirmed by genotyping PCR.

### Generation of chimaeras

#### Human-mouse chimaeras

hPSCs were trypsinized and suspended in their culture medium supplemented with 2 µM Y. 4–10 hPSCs were injected into the perivitelline space of 8-cell/morulae-stages embryos using a piezo-driven or laser micro-manipulator. The injected embryos were cultured in KOSM medium containing 30 %(v/v) StemFlex and 2 µM Y, covered with mineral oil. By the following day, the injected embryos developed to the early blastocyst stage. These blastocysts were transferred into the uteri of pseudo-pregnant recipient ICR mice at 2.5 dpc. For transgene induction, drinking water containing 2 mg/mL Dox and 35 mg/mL sucrose was administered to the recipient mice starting from the day of embryo transfer.

#### Human-pig chimaeras

10-20 hPSCs were transferred into a drop of N2B27 medium containing 2 µM Y and 10-50 µg/mL PHA-P, then injected into the blastocoel cavity adjacent to the ICM of early blastocyst (E5) stage embryos using piezo-driven or laser micromanipulator. Injected embryos were cultured in N2B27 medium supplemented with 2 µM Y for one day and 20 µg/mL Dox was added following day and fixed the embryos at E7 for preimplantation embryo analysis. For post implantation embryo analysis, the injected embryos were cultured N2B27 medium supplemented with 2 µM Y for one day and transferred into the uteri of recipient gilts at E6. The recipient gilts were fed 50 mg/kg/day Dox-containing food starting 3 days before embryo transfer and continuing until the day before or on the day of sampling. Recipient gilts were euthanized, and embryos were collected from the uteri by flushing with PBS containing 1% foetal calf serum.

### Analysis of interspecies chimaeras

#### Chimerism estimation by ddPCR

Human cell frequency was estimated by quantifying the copy number of human mitochondrial DNA relative to the total mammalian genome in chimeric tissues. Absolute copy numbers were measured using the QX200 Droplet Digital PCR System (Bio-Rad) with ddPCR Supermix for Probes (no dUTP) (Bio-Rad, Cat# 1863024) following the manufacturer’s instructions. Tissues were homogenized using the TissueLyser LT (Qiagen, Cat# 85600) in ProK lysis buffer (20 mM Tris-HCl (pH 8.0), 100 mM NaCl, 5 mM EDTA, 0.1% SDS, 200 µg/mL Proteinase K). Genomic DNA was extracted using the NucleoSpin Tissue Kit (MACHEREY-NAGEL, Cat# 740952.250). DNA from donor hPSCs was used as a reference. For each 20 µL ddPCR reaction, 30–100 μg of genomic DNA was digested with HindIII-HF (New England Biolabs, Cat# R3104S). PCR was performed using the following primers and probes (IDT): human mitochondria ddPCR primer F 5’-CGTACGCCTAACCGCTAACA-3’; R 5’-GATAATGCTAGGGTGGCGCT-3’, human mitochondria ddPCR probe 5’-6-FAM/TGCAGGCCA/ZEN/CCTACTCATGCA/IABkFQ-3’. Zeb2 ddPCR primer F 5’-GGATGGGGAATGCAGCTCTT-3’; R 5’-AGTGCGGCAGAATACAGCA-3’, Zeb2 ddPCR probe 5’-HEX/TGATGGGTT/ZEN/GTGAAGGCAGCTGCACCT/IABkFQ-3’. The Zeb2 primers and probe are cross-reactive with mouse and human. Chimaerism was calculated as the number of mitochondrial copies of each embryo divided by the Zeb2 gene copy number and normalized by the average number of mitochondria in the donor hPSCs. Data analysis was performed using QX Manager v1.1 (Bio-Rad).

#### FACS and Genotyping

Hearts were isolated from E12.5 chimeric embryos, minced, and dissociated with 0.25% Trypsin-EDTA for 30 min at 37°C. Dissociated cells were sorted into EGFP-positive and EGFP-negative fractions using an SH800 cell sorter (Sony), and 1000 cells were collected per fraction. PB001 hPSCs and ICR mouse skin cells were processed in the similar manner. The samples were Genomic DNA was extracted using the Proteinase K method, and PCR was performed with Tks-Gflex (Takara) and primers: human β-actin (ACTB) primer F 5’-TGACATGGTGTATCTCTGCCT-3’; mouse ACTB primer F 5’-GCTCTTTCCCAGACGAGGTCT-3’; human-mouse ACTB common primer R 5’-CAGGAGGAGCAATGATCTGA-3’. Chimerism in the heart was quantified based on EGFP signal using FlowJo v10.9 software (BD Biosciences).

### Immunohistochemistry

#### Preimplantation embryos

Embryos were fixed in 4% paraformaldehyde (PFA) in PBS for 15 min at room temperature. Samples were rinsed in PBS containing 3 mg/mL polyvinylpyrrolidone (PBS/PVP) and permeabilised with PBS/PVP containing 0.25% Triton X-100 for 30 min. Blocking was performed in embryo blocking buffer (PBS supplemented with 0.1% BSA, 0.01% Tween-20 and 2% donkey serum) for 2–3 h at 4°C. Embryos were incubated with primary antibody (1:100 dilution) in blocking buffer overnight at 4°C, followed by three times washes in blocking buffer and incubation with secondary antibody (1:200 dilution) in blocking buffer containing 500 ng/mL DAPI for 1–2 h at room temperature in the dark. Samples were then rinsed three times for 15 min each in blocking buffer.

#### E6.5 embryos

Embryos were dissected from the uterus under a stereo-microscope. Embryos were fixed in 4% PFA for 1 h at 4°C and washed three times in PBS. Samples were permeabilised in 0.5% Triton X-100/PBS for 1 h at room temperature, then blocked with MaxBlock Blocking Medium for 1 h at 37°C. After three PBS washes, embryos were incubated with primary antibodies diluted in 0.5% Triton X-100/PBS overnight at 4°C. Following three PBS washes, sample were incubated with secondary antibodies (1:200 dilution) in 0.5% Triton X-100/PBS for 1–2 h at room temperature in the dark. Samples were washed three times with PBS and cleared overnight at room temperature in CUBIC1 solution containing 500 ng/mL DAPI.

#### Embryonic and extraembryonic tissues

Embryonic and extraembryonic tissues were dissected from the uterus under a stereo-microscope. Tissues were fixed in 4% PFA for 24 h at 4°C, followed by three washes in PBS. For CD31 staining, fresh tissues were incubated with a fluorophore-conjugated anti-CD31 antibody in PBS for 12–24 h at 4°C and fixed in 4% PFA for 12–24 h at 4°C. Samples were permeabilised in 0.5% Triton X-100/PBS for 2 h at room temperature and blocked with MaxBlock Blocking Medium for 1 h at 37°C. After three PBS washes, samples were incubated with fluorophore-conjugated antibodies in 0.5% Triton X-100/PBS for 24 h at 4°C. Following three PBS washes, tissues were cleared overnight at room temperature in CUBIC1 containing 500 ng/mL DAPI.

All antibodies are listed in the Key Resources Table.

### Culture of hPSCs-derived cardiomyocytes

Hearts were isolated from E13.5 chimaeric embryos, minced, and dissociated with 0.25% Trypsin-EDTA for 30 min at 37°C. EGFP-positive cells were sorted using an SH800 cell, seeded on MEFs, and cultured in MEF medium for two days at 37°C with 5% CO_2_.

### Imaging

Immunostained samples were imaged using confocal laser scanning microscopes FV3000 (Olympus) or SP8 (Leica). Dissected embryos and organs were imaged using a fluorescent stereo microscope M164FC (Leica). Live imaging of cells was performed using BZ-X810 (Keyence).

### Single-cell RNA-seq of chimeric embryos

The SMART-Seq HT (Takara Bio Inc., Cat# 634438) was used following the manufactures’ instructions with minimal modifications to synthesize cDNA from single cells of chimeric embryos. E6.5 chimaeric embryos were dissected and the embryonic region was further micro-dissected using fine glass capillaries or needles in PBS with 0.1% BSA and 10 µM Y. Individual epiblast regions were incubated in dissociation buffer (PBS containing 0.05% trypsin, 1.2 mM EDTA, 5 U/mL Dispase ll and 0.1% BSA) at 37°C in a CO_2_ incubator for 5 min. The embryos were dissociated by gently pipetting and transferred to a drop of medium on a dish. E12.5 chimaeric hearts and intestines were dissected, trimmed and dissociated with 0.25% Trypsin-EDTA for 30 min ay 37°C. Individual GFP+ human cells were manually collected under a microscope using micro-manipulators and transferred into PCR tubes prefilled with 3 µL cold nuclease-free PBS. A mixture of 6 µL of condensed lysis buffer (0.76 µL 10× lysis buffer, 0.04 µL RNase inhibitor, 0.8 µL 3′SMART-Seq CDS Primer II A, and 4.4 µL nuclease-free water) was added, pipetted several times to mix, snap-frozen on dry ice, and stored at -80°C until cDNA synthesis was performed. 10 µL of one-step master mix (0.56 µL nuclease-free water, 6.4 µL one-step buffer, 0.8 µL SMART-Seq HT oligonucleotide, 0.4 µL RNase inhibitor, 0.24 µL SeqAmp DNA polymerase, and 1.6 µL SMARTScribe Reverse Transcriptase) was added to 9-10 µL of the lysate and subjected to reverse transcription followed by 20 cycles cDNA amplification in a thermal cycler (TP350, Takara). cDNA libraries were purified with SPRIselect beads (Beckman Coulter, Cat# B23318) or AMPure XP beads (Beckman Coulter, Cat#A63881) and quantified by Qubit 3 fluorometer (Thermo, Cat# Q33216) or Quant-iT PicoGreen dsDNA Assay Kit (Thermo Fisher Scientific Inc., Cat#P11496). Illumina libraries were prepared using Nextera XT (Illumina, Cat# FC-131-1024) according to the manufactures’ instructions and sequenced on HiSeq X (150 bp paired-end), NovaSeq (150 bp paired-end), NovaSeq X plus (150 bp paired-end), or NextSeq (75 bp paired-end). The fastq files were adaptor-trimmed by Trimmomatic (Galaxy v0.38.1) and mapped to hg38 and mm10 reference genomes using HISAT2 (Galaxy v2.1.0+galaxy5) on Galaxy (v20.05). The BAM files were subjected to XenofilteR (v1.6) on R (v4.2.2) for computational deconvolution of mouse and human reads with MM-threshold 4 for PE75 or 8 for PE150. Species-specific BAM files were counted by featureCounts (Galaxy v 1.6.4+galaxy2) based on the GTF file of Ensemble, version-release.98. Transcript integrity number (TIN) and Gene Body Coverage were analysed by RSeQC (Galaxy v2.6.4.1 and Galaxy v2.6.4.3) to exclude low-quality samples. The count was normalized by DESeq2 (v1.38.2) and analysed for differentially expressed genes (DEGs).

### Bulk RNA-seq of chimaeric embryos

E13.5 chimaeric embryos were dissected, and GFP+ organ pieces were trimmed under fluorescence stereoscopy (Leica, Cat# M165FC). Total RNA was purified using the RNeasy Mini (Qiagen, Cat# 74134) after tissue homogenization with TissueLyser LT in TRIzol (Thermo, Cat# 15596-018). cDNA libraries were generated from 5 ng total RNA in 20 µL reactions and sequentially amplified by 11 cycles by SMART-seq HT. Illumina libraries were prepared by Nextera XT following the manufacturer’s instructions and sequenced using NovaSeq (150 bp paired-end). Data were processed using the same pipeline as described above for scRNA-seq.

### Statistics

Graphs show mean ± standard deviation (SD). Statistical significance was assessed using unpaired Student’s t-test, except for comparisons marked with †, which were analyzed using Welch’s t-test. n.s., not statistically significant; *p < 0.05, **p < 0.01.

