## Supplementary_Figures for "The generation of viable, structurally integrated human-mouse chimaeras through enhanced hPSCs proliferation"

### Supplementary Figure 1

A

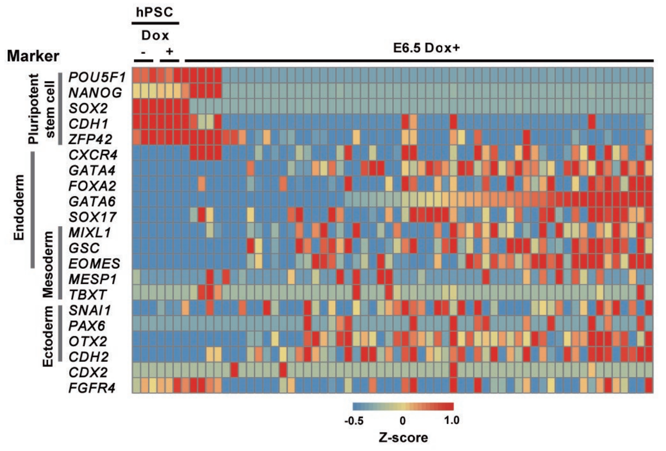


B

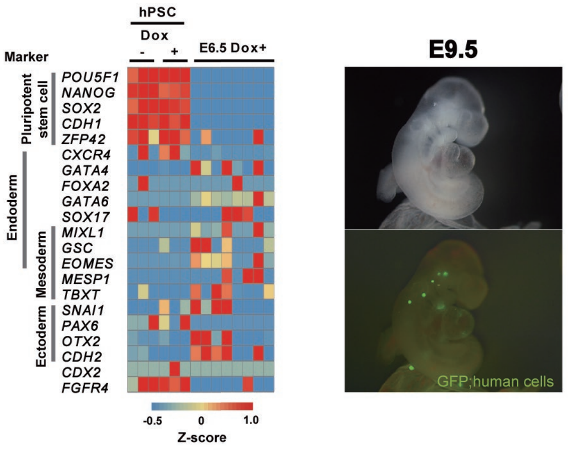


C

| Donor cell name | Trans gene | | Number of injected cells | Host embryo strain | Dox treated | Developmental stages | Transferred embryos | Implantation[a] | live fetus[b] (b/a %) | Chimeras[c] (c/a %) |
|---|---|---|---|---|---|---|---|---|---|---|
| PB001#4 | CAG-BCL2 | - | 5-8 | BDF1xB6 | - | E13.5 | 41 | 16 | 12 (75%) | 3 (19%) |
| PB001#4 | CAG-BCL2 | TRE-MYCL | 5-8 | BDF1xB6 | E2.5 to E13.5 | E13.5 | 20 | 15 | 6 (40%) | 5 (33%) |
| PB001#4 | CAG-BCL2 | TRE-MYCN | 5-8 | BDF1xB6 | E2.5 to E13.5 | E13.5 | 16 | 16 | 11 (69%) | 6 (38%) |
| PB001#4 | CAG-BCL2 | TRE-MYCN | 5-8 | BDF1xB6 | E2.5 to E8.5 | E13.5 | 10 | 5 | 4 (75%) | 3 (60%) |
| PB001#4 | CAG-BCL2 | TRE-MYC | 5-8 | BDF1xB6 | E2.5 to E13.5 | E13.5 | 32 | 28 | 5 (18%) | 4 (14%) |
| PB001#4 | CAG-BCL2 | TRE-MYC | 5-8 | BDF1xB6 | E2.5 to E8.5 | E13.5 | 36 | 25 | 6 (24%) | 5 (20%) |
| PB001#4 | CAG-BCL2 | - | 5-8 | BDF1xB6 | - | E16.5 | 36 | 33 | 8 (24%) | 2 (6%) |
| PB001#4 | CAG-BCL2 | TRE-MYCL | 5-8 | BDF1xB6 | E2.5 to E16.5 | E16.5 | 30 | 19 | 4 (21%) | 3 (16%) |
| PB001#4 | CAG-BCL2 | TRE-MYCN | 5-8 | BDF1xB6 | E2.5 to E16.5 | E16.5 | 24 | 22 | 16 (73%) | 8 (36%) |
| PB001#4 | CAG-BCL2 | TRE-MYCN | 5-8 | BDF1xB6 | E2.5 to E8.5 | E16.5 | 54 | 29 | 17 (59%) | 5 (9%) |

D

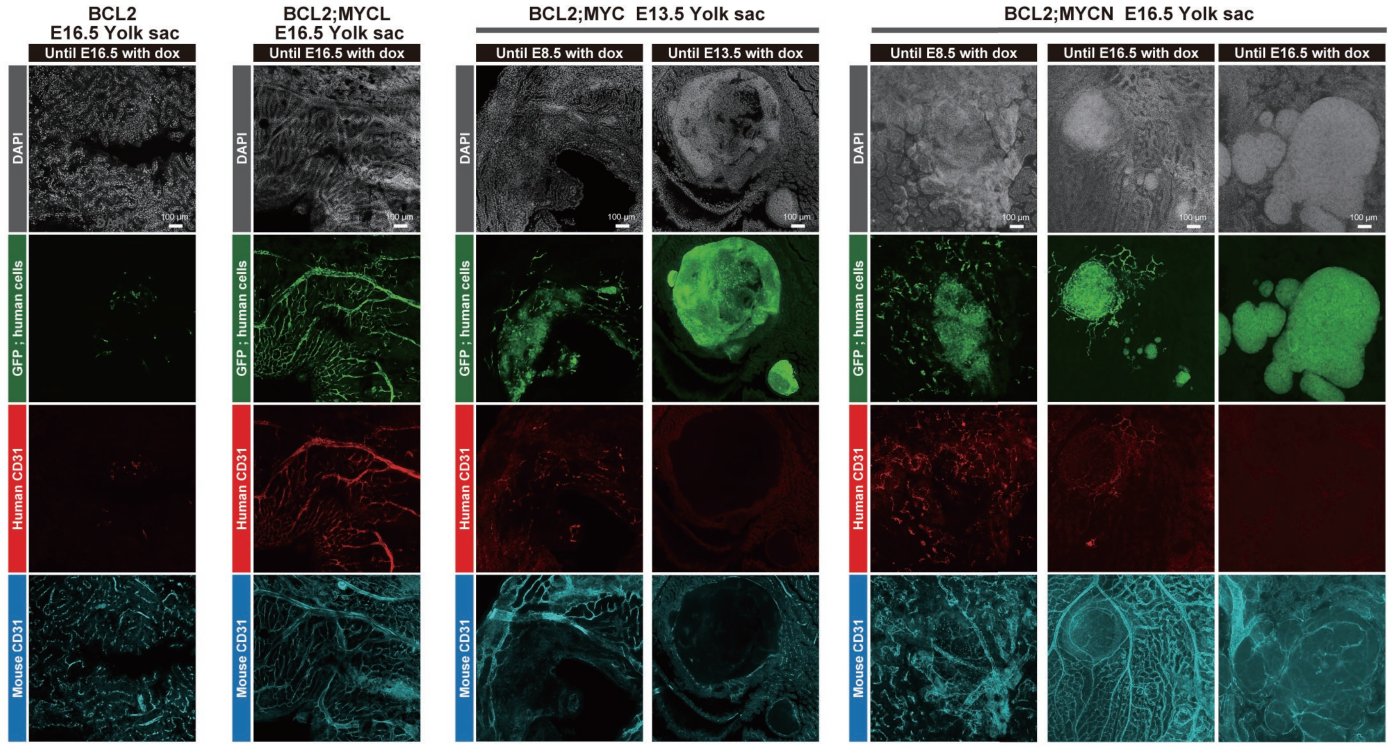


E

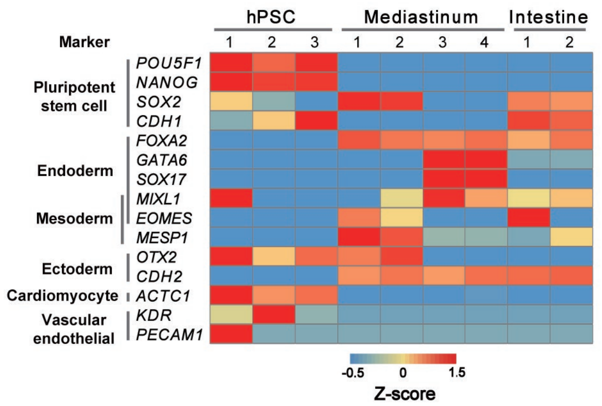


F

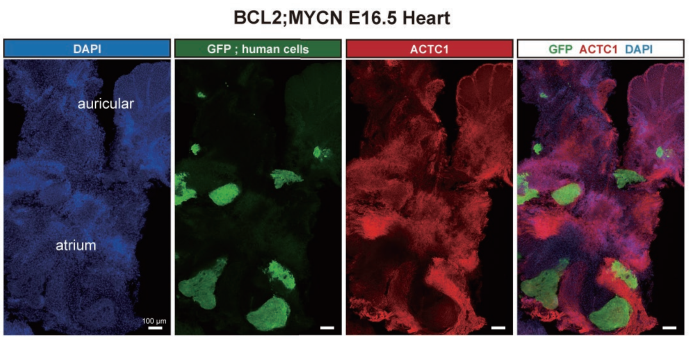

### Supplementary Figure2

**A**

| Donor cell name | Trans gene | | Nunber of injected cells | Host embryo strain | Dox treated | Developmental stages | Transferred embryos | Implantation[a] | live fetus[b] (b/a %) | Chimeras[c] (c/a %) |
|---|---|---|---|---|---|---|---|---|---|---|
| PB001#4 | CAG-BCL2 | - | 5-8 | ICR | E2.5 to E12.5 | E12.5 | 56 | 34 | 20(59%) | 9(26%) |
| PB001#4 | CAG-BCL2 | TRE-CDK9 | 5-8 | ICR | E2.5 to E12.5 | E12.5 | 56 | 38 | 17(44%) | 7(18%) |
| PB001#4 | CAG-BCL2 | TRE-E2F1 | 5-8 | ICR | E2.5 to E12.5 | E12.5 | 52 | 43 | 22(51%) | 0(0%) |
| PB001#4 | CAG-BCL2 | TRE-E2F2 | 5-8 | ICR | E2.5 to E12.5 | E12.5 | 77 | 48 | 27(56%) | 13(27%) |
| PB001#4 | CAG-BCL2 | TRE-E2F3a | 5-8 | ICR | E2.5 to E12.5 | E12.5 | 80 | 43 | 31(72%) | 10(23%) |
| PB001#4 | CAG-BCL2 | TRE-E2F4 | 5-8 | ICR | E2.5 to E12.5 | E12.5 | 36 | 34 | 15(44%) | 8(24%) |
| PB001#4 | CAG-BCL2 | TRE-MYCL | 5-8 | ICR | E2.5 to E12.5 | E12.5 | 48 | 42 | 22(52%) | 12(29%) |

**B**

| Donor cell name | Trans gene | | Nunber of injected cells | Host embryo strain | Dox treated | Developmental stages | Transferred embryos | Implantation[a] | live fetus[b] (b/a %) | Chimeras[c] (c/a %) |
|---|---|---|---|---|---|---|---|---|---|---|
| 7F 3669#1 | CAG-BCL2L1 | TRE-MYCL | 5-8 | BDF1xB6 | E2.5 to E12.5 | E12.5 | 44 | 23 | 8(35%) | 6(26%) |
| 7F 3669#1 | CAG-BCL2L1 | TRE-MYCL | 5-8 | ICR | E2.5 to E12.5 | E12.5 | 59 | 17 | 8(47%) | 5(29%) |

**C**

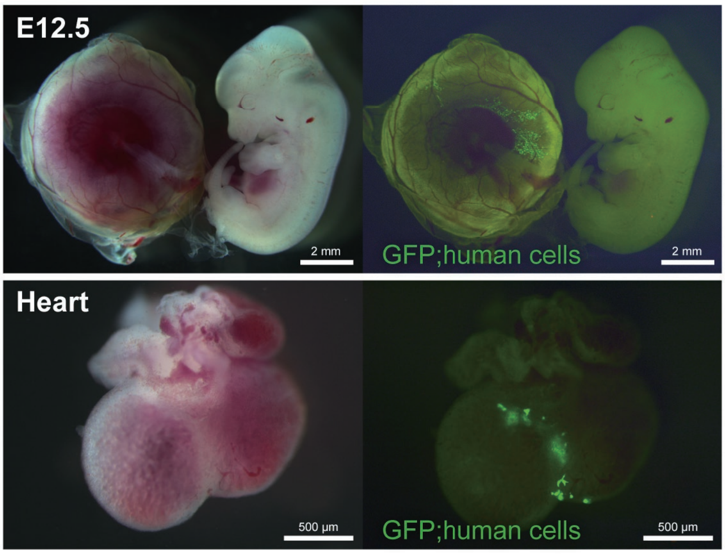


**D**

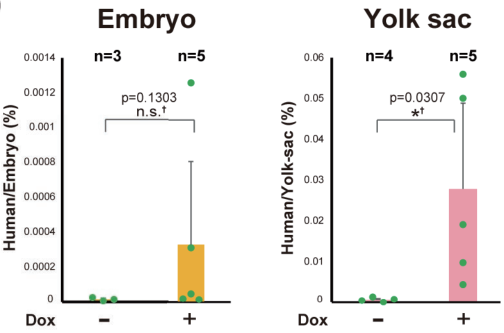


| Dox | − | + |
|---|---|---|
| Aberage | $1.50 \times 10^{-5}$ | $3.26 \times 10^{-4}$ |
| SD | $7.80 \times 10^{-6}$ | $4.76 \times 10^{-4}$ |
| Max | $2.51 \times 10^{-5}$ | $1.25 \times 10^{-3}$ |

| Dox | − | + |
|---|---|---|
| Aberage | $6.29 \times 10^{-4}$ | $2.78 \times 10^{-2}$ |
| SD | $4.89 \times 10^{-4}$ | $2.11 \times 10^{-2}$ |
| Max | $1.35 \times 10^{-3}$ | $5.59 \times 10^{-2}$ |

### Supplementary Figure3

**A**

| Donor cell name | Trans gene | | Nunber of injected cells | Host embryo strain | Dox treated | Developmental stages | Transferred embryos | Implantation[a] | live fetus[b] (b/a %) | Chimeras[c] (c/a %) |
|---|---|---|---|---|---|---|---|---|---|---|
| PB001#4 | CAG-BCL2 | TRE-MYCL | 6-10 | BDF1 x B6 | - | E9.5 | 20 | 5 | 5(100%) | 5(100%) |
| PB001#4 | CAG-BCL2 | TRE-MYCL | 6-10 | BDF1 x B6 | - | E12.5 | 60 | 38 | 7(18%) | 7(18%) |
| PB001#4 | CAG-BCL2 | TRE-MYCL | 6-10 | BDF1 x B6 | E2.5 to E9.5 | E9.5 | 51 | 30 | 13(43%) | 6(20%) |
| PB001#4 | CAG-BCL2 | TRE-MYCL | 6-10 | BDF1 x B6 | E2.5 to E12.5 | E12.5 | 70 | 59 | 26(44%) | 17(29%) |
| PB001#4 | CAG-BCL2 | TRE-MYCL | 4-5 | BDF1 x B6 | - | E12.5 | 40 | 38 | 22(58%) | 5(13%) |
| PB001#4 | CAG-BCL2 | TRE-MYCL | 4-5 | BDF1 x B6 | E2.5 to E6.5 | E12.5 | 40 | 26 | 18(69%) | 11(42%) |
| PB001#4 | CAG-BCL2 | TRE-MYCL | 4-5 | BDF1 x B6 | E2.5 to E9.5 | E12.5 | 36 | 18 | 9(50%) | 5(28%) |
| PB001#4 | CAG-BCL2 | TRE-MYCL | 4-5 | BDF1 x B6 | E2.5 to E12.5 | E12.5 | 68 | 46 | 13(28%) | 7(15%) |

**B**

**E12.5**
**Heart**

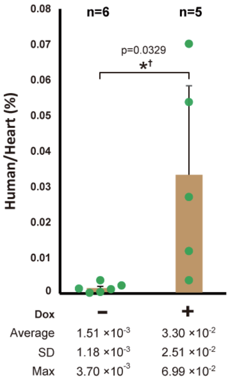


**C**

**E12.5**
**Pooled heart samples**

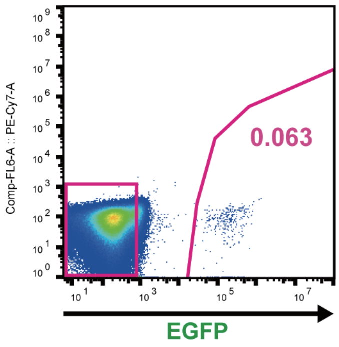


**D**

**Genomic PCR**

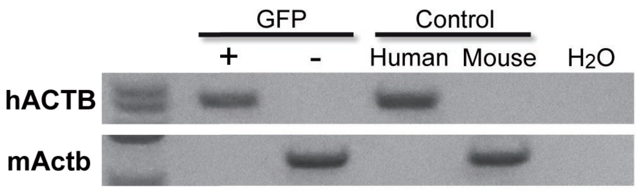


**E**

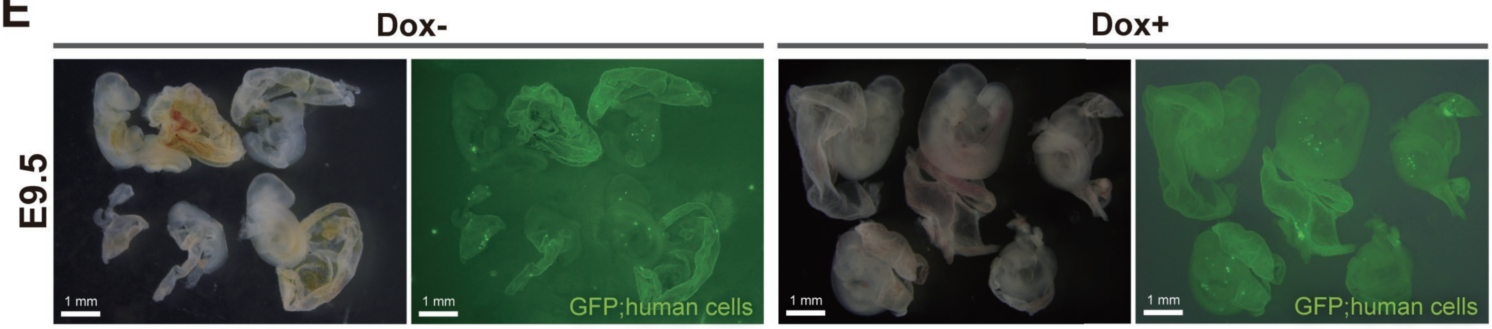

### Supplementary Figure4

## A

| Donor cell name | Trans gene | | Nunber of injected cells | Host embryo strain | Dox treated | Developmental stages | Transferred embryos | Implantation[a] | live fetus[b] (b/a %) | Chimeras[c] (c/a %) |
|---|---|---|---|---|---|---|---|---|---|---|
| PB001#4 | TRE-BCL2L1 | TRE-MYCL | 4-7 | BDF1xB6 | E2.5 to E9.5 | E19.5 | 61 | - | 22 | 2 |
| PB001#4 | CAG-BCL2 | TRE-MYCL | 4-5 | BDF1xB6 | E2.5 to E19.5 | E19.5 | 32 | 13 | 9 (69%) | 4 (31%) |

## B

TRE-BCL2L1 TRE-MYCL

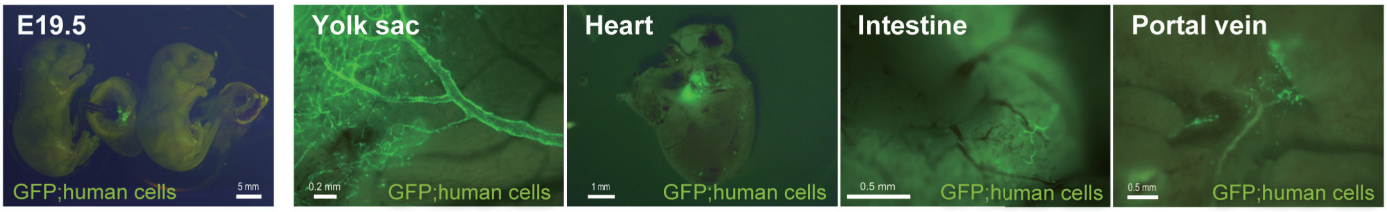


## C

CAG-BCL2 TRE-MYCL E19.5 Brain

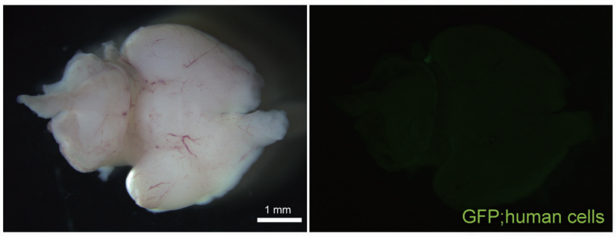


## D

| Donor cell name | Trans gene | | Nunber of injected cells | Host embryo strain | Dox treated | Developmental stages | Transferred embryos | Implantation[(a)] | live fetus[(b)] (b/a %) | Chimeras[(c)] (c/a %) |
|---|---|---|---|---|---|---|---|---|---|---|
| PB001#4 | CAG-BCL2 | TRE-MYCL | 5-8 | BDF1xB6 | - | E6.5 | 44 | 20 | 18(90%) | 17(80%) |
| PB001#4 | CAG-BCL2 | TRE-MYCL | 5-8 | BDF1xB6 | E2.5 to E6.5 | E6.5 | 38 | 34 | 26(76%) | 26(76%) |

## E

E2.5 mouse embryo + CAG-BCL2 TRE-MYCL hPSCs, 1-day culture

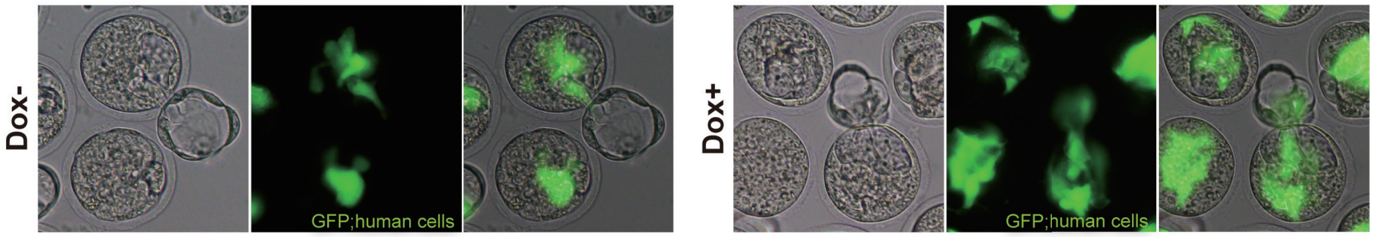


## F

CAG-BCL2 TRE-MYCL E6.5

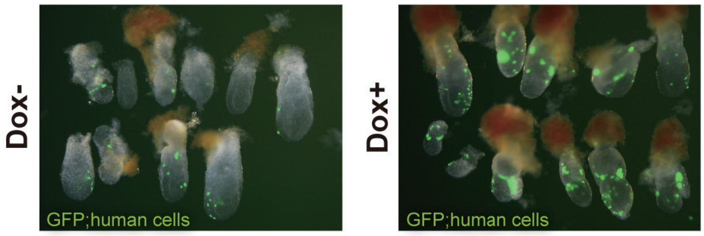


## G

| Donor cell name | Trans gene | | Nunber of injected cells | Host embryo strain | Dox treated | Developmental stages | Transferred embryos | Implantation | live fetus | Chimeras |
|---|---|---|---|---|---|---|---|---|---|---|
| PB001#4 | CAG-BCL2 | TRE-MYCL | 10-20 | Pig | + | E22 | 61 | - | 21 | 3 |
| PB001#4 | CAG-BCL2 | TRE-MYCL | 10-20 | Pig | + | E22 | 58 | - | 40 | 3 |
