## Supplementary_Figure_Legends for "The generation of viable, structurally integrated human-mouse chimaeras through enhanced hPSCs proliferation"

**SUPPLEMENTAL FIGURE LEGENDS**

**Supplementary Table 1.**
RNA-seq count matrix of donor hPSCs and hPSCs derivatives isolated from E6.5 human-mouse chimaeras in this study.

**Supplementary Video 1.**

Spontaneous beating of *BCL2*;*MYCL*-hPSC derivatives isolated from E13.5 human-mouse chimaeric hearts. Related to Figure 3.

**Figure S1 Related to Figure 1**

(A) Heatmap of gene expression profiles from scRNA-seq of undifferentiated hPSCs and *BCL2*-hPSC derivatives isolated from E6.5 human-mouse chimaeras.
(B) Heatmap of gene expression profiles from scRNA-seq of undifferentiated hPSCs and *BCL2*;*MYC*-hPSC derivatives isolated from E6.5 human-mouse chimaeras, and representative images of E9.5 human-mouse chimaeras derived from *BCL2*;*MYCL*-hPSCs.
(C) Summary of embryo manipulation results for the generation of human-mouse chimaeras corresponding to Figure 1.
(D) Single-channel fluorescence images of yolk sacs from E13.5 or E16.5 chimaeras derived from GFP-labelled *BCL2*-, *BCL2*;*MYC*-, *BCL2*;*MYCN*- or *BCL2*;*MYCL*-hPSCs under varying duration of Dox administration corresponding to Fig.1B.
(E) Heatmap of bulk RNA-seq gene expression profiles in hPSCs and *BCL2*;*MYC*-hPSC-derived cell clusters isolated from the mediastinum and intestine of *BCL2*;*MYC* human-mouse chimaeras.
(F) Representative immunohistochemical staining images of E16.5 hearts from *BCL2*;*MYCN*- human-mouse interspecies chimaera.

**Figure S2. Related to Figure 2**

(A) Summary of embryo manipulation results for the generation of human-mouse chimaeras corresponding to Figure 2.
(B) Summary of embryo manipulation results for BCL2;MYCL human-mouse chimaeras　derived from a different hPSC line.
(C) Representative images of E12.5 human-mouse chimaeras and a dissected heart derived from GFP-labelled 7F 3669#1 *BCL2*;*MYCL*-hPSCs.
(D) ddPCR analysis of human cell chimaerism in whole embryos and yolk sacs from E12.5 chimaeras. Each dot represents one chimaeric embryos.

Graphs show mean ± standard deviation (SD). Statistical significance was assessed using unpaired Student’s t-test, except for comparisons marked with †, which were analyzed using Welch’s t-test. n.s., not statistically significant; *p < 0.05, **p < 0.01.

**Figure S3. Related to Figure 3**

(A) Summary of embryo manipulation results for the generation of human-mouse chimaeras corresponding to Figure 3.
(B) ddPCR analysis of human cell chimaerism in isolated hearts from E12.5 human-mouse chimaeras derived from GFP-labelled *BCL2*;*MYCL*-hPSCs. Each dot represents one embryo or tissue sample.
(C) Flow cytometry analysis of E12.5 hearts from GFP-labelled *BCL2*;*MYCL* human-mouse chimaeras. Dox was administer until the time of dissection.
(D) Genomic PCR on EGFP^+^ and EGFP^–^ cells isolated from E12.5 hearts of human-mouse chimaeras. Control human cells; PB001 hPSC, control mouse cells; ICR mouse.
(E) Representative images of E9.5 *BCL2*;*MYCL* human-mouse chimaeras with or without Dox administration, corresponding to Figure 3D.

Graphs show mean ± standard deviation (SD). Statistical significance was assessed using unpaired Student’s t-test, except for comparisons marked with †, which were analyzed using Welch’s t-test. n.s., not statistically significant; *p < 0.05, **p < 0.01.

**Figure S4. Related to Figure 4**

(A) Summary of embryo manipulation results for the generation of human-mouse chimaeras corresponding to Figures 4A and S4B-C.
(B) Representative images of E19.5 human-mouse chimaeras derived from GFP-labelled *BCL2L1*;*MYCL*-hPSCs. Dox was administered until E9.5.
(C) Representative images of the brain from E19.5 *BCL2*;*MYCL* human-mouse chimaeras. Dox was administered until the time of dissection.
(D) Summary of embryo manipulation results for the generation of human-mouse chimaeras corresponding to Figure 4B-C.
(E) Representative images of E3.5 human-mouse chimaeras derived from GFP-labelled *BCL2*;*MYCL*-hPSCs.
(F) Representative immunohistochemical staining images of E6.5 embryos from *BCL2*;*MYCL* human-mouse chimaera.
(G) Summary of embryo manipulation results for the generation of human-pig chimaeras corresponding to Figure 4D-F.
